# Protein-mediated stabilization and nicking of the non-template DNA strand dramatically affect R-loop formation *in vitro*

**DOI:** 10.1101/2025.04.24.649451

**Authors:** Ethan Holleman, Thomas E. Catley, Tadas Sereiva, Stella R. Hartono, Alice L.B. Pyne, Frédéric Chédin

## Abstract

R-loops are an important class of non-B DNA structures that form co-transcriptionally. Using *in vitro* transcription and unbiased quantitative sequencing readouts, we show that the addition of single-strand DNA binding proteins co-transcriptionally can drive a 3- to 5-fold increase of R-loop frequency without significant changes to R-loop distribution. We propose that this is caused by stabilizing and preventing the collapse of short nascent R-loops. This suggests that R-loop formation is highly dynamic and highlights single strand binding proteins as players in cellular R-loop regulation. We further show that non-template strand DNA nicks are powerful initiators of R-loop formation, increasing R-loop frequencies by up to two orders of magnitude. Atomic force microscopy (AFM) revealed that the non-template strand in nick-initiated structures is often flayed away from the RNA:DNA hybrid and engaged in self-pairing, creating unique forked R-loop features. DNA nicks, one of the most frequent DNA lesions in cells, are therefore potential hotspots for opportunistic R-loop initiation and may cause the formation of a novel class of R-loops. Overall, this work highlights the importance of the displaced single-strand on R-loop initiation and dynamics.

**SIGNIFICANCE STATEMENT:** R-loops are three-stranded DNA:RNA hybrid structures that form during transcription and play critical roles in both gene regulation and genome stability. This study reveals that single-stranded DNA-binding protein (SSB) increases R-loop frequency by stabilizing nascent R-loops, revealing a role for this protein in R-loop dynamics. Additionally, single strand DNA breaks, common cellular lesions, are identified as potent initiators of R-loops, increasing their rate of formation by up to 100-fold. Using atomic force microscopy, the study uncovers unique forked structures in nick-initiated R-loops, representing a novel class of secondary R-loop structures. These findings provide new insights into how cellular factors and DNA damage influence R-loop formation, with implications for understanding genome instability in health and disease.

## INTRODUCTION

R-loops are three-stranded nucleic acid structures that form during transcription upon reannealing of the nascent RNA to the template DNA strand. These structures have been described from bacteria, yeasts, plants, metazoans, and mammals (1) where they form prevalently over transcribed regions. Under physiological conditions, R-loops are well-tolerated and associated with a number of adaptive processes including transcription termination (2),(3) chromatin patterning (4), class switch recombination (5) and the regulation of gene expression (6). At the same time, harmful R-loops have been implicated as players in a variety of maladaptive cellular processes, in particular as threats to genomic stability (6–8), and a number of human diseases have been linked to deregulated R-loop metabolism (9). Despite the growing recognition of the complex roles played by R-loops, significant gaps remain in our understanding of the fundamental principles guiding R-loops initiation, stability, and decay. Biochemical reconstitution of R-loop formation using *in vitro* transcription studies combined with mathematical modeling have identified DNA sequence and DNA topology as key factors regulating R-loop formation. R-loops are favored over GC-rich and GC-skewed regions owing to the relative energetic favorability of the resulting RNA:DNA hybrids over the corresponding DNA duplex (10). R-loops are also favored on negatively supercoiled DNA templates (11–13) owing to their ability to absorb negative superhelicity locally and to relax the surrounding DNA fiber to a lower energetic state (12). Recent mathematical modeling of R-loop energetics at equilibrium revealed that R-loop formation becomes favorable when the combined energetic return due to DNA sequence and DNA topology is such that the energetic barrier caused by the formation of Y-junctions at the start and end of R-loops can be overcome (12). It is likely, however, that additional factors beyond DNA sequence and topology play important, yet-to-be understood, roles in driving R-loop formation or stability.

The impact of the non-template DNA strand on R-loop formation, dynamics, and stability, has not been fully considered. This strand is in competition with the nascent RNA for reannealing to the template DNA strand (14, 15). DNA reannealing will affect the ability of an R-loop to initiate by preventing the formation of the RNA:DNA hybrid, or, if reannealing occurs after R-loop initiation, it could cause the collapse of nascent R-loops and the displacement of the RNA. This competition model predicts that stabilization of the looped out non-template strand may reduce R-loop collapse. Several studies have suggested that G quadruplex (G4) formation on the G-rich displaced strand stabilizes R-loops due to the fact that that strand is now self-paired and additional energy is required to melt it back to ssDNA before it could reanneal (16) (17). Binding of single-stranded DNA binding proteins on the displaced strand may play a similar role, although this has not been formally assessed in *in vitro* transcription studies. Replication Protein A (RPA) was reported to catalyze R-loop formation *in vitro* (18) and to bind to R-loops *in vivo*, facilitating the recruitment of the R-loop resolving enzyme Ribonuclease H1 (RNase H1) (19). However, its role during co-transcriptional R-loop formation has not been measured. It has also been reported that single-stranded DNA breaks (nicks) in the non-template strand can stimulate R-loop formation (20). This effect was attributed to an increased ability of the non-template strand downstream of the nick to dissociate from the template DNA strand, perhaps through local breathing, facilitating RNA invasion and RNA:DNA hybrid formation. However, the positions, lengths, or characteristics of the R-loops associated with nicks were not determined, and the impact of nicks on R-loop formation frequency was not quantitatively assessed.

Here, we used *in vitro* transcription assays on defined plasmid substrates followed by single-molecule R-loop footprinting (SMRF-seq (13)) to quantitatively measure the impact of single-stranded DNA binding proteins and non-template DNA nicks on R-loop formation. This work reveals that each of these factors exerts a profound effect on the R-loop landscape that is often equally or more powerful than the well-documented effect of DNA sequence and DNA topology.

## RESULTS

### Single strand binding proteins stabilize nascent R-loop formation

In an R-loop, the RNA strand of the RNA:DNA hybrid and the non-template DNA strand compete for binding to the template DNA strand. If the DNA strand prevails, the R-loop will not form or it will collapse post initiation. Here, we tested the hypothesis that single-strand DNA binding proteins may favor R-loop formation by stabilizing the non-template DNA strand. To test this, we used *in vitro* transcription (IVT) reactions with the T7 RNA polymerase on various R-loop prone templates, focusing first on the effect of the *E. coli* SSB protein. Under co-transcriptional conditions (co-txn), SSB was added to the reaction prior to transcription so that SSB may interact with R-loops from the moment they initiate. As a control, we first performed IVT, and SSB was only added after the transcription was terminated, such that SSB may only interact with R-loops that exist at the end of the reaction (post-txn).

Co-transcriptional SSB addition led to a slight but discernible RNase H-sensitive mobility shift on agarose gels that was not seen upon addition of SSB post-transcription (**Fig S1A**). To quantitatively measure R-loop frequencies and positions in an unbiased manner, we performed SMRF-seq after IVT. Three different linear plasmid constructs, each with a different background propensity for R-loop formation based on DNA sequence, were used. Overall, adding SSB co-transcriptionally led to a remarkable 3- to 5-fold increase in R-loop frequencies compared to SSB addition post-transcription (**Fig 1A,B**). As expected, R-loop peaks measured by SMRF-seq were highly-strand specific and located to the non-template strand. R-loop footprints were also highly sensitive to RNase H pre-treatment prior to SMRF-seq (**Fig S1B**). Interestingly, R-loop locations did not significantly change upon SSB addition (**Fig 1A, S1C**). These observations are most consistent with the notion that SSB binds to the exposed non-template strand of an R-loop once it becomes available after an R-loop has initiated. SSB binding is proposed to stabilize that strand and lower the chances of early R-loop dissolution by spontaneous DNA reannealing. The strong effect of SSB on R-loop frequencies indicates that, in the absence of SSB, the large majority of R-loops collapse before they can be measured. Interestingly, it should be noted that all samples here were subjected to de-proteinization prior to SMRF-seq. This suggests that the increase in R-loops observed upon co-transcriptional SSB addition was derived from a population of R-loops that were prone to early collapse without SSB, but did not require SSB for their long-term stability once fully formed. Thus, the effect of SSB is most likely felt early during R-loop formation through the stabilization of short nascent R-loops.

**Figure 1.**
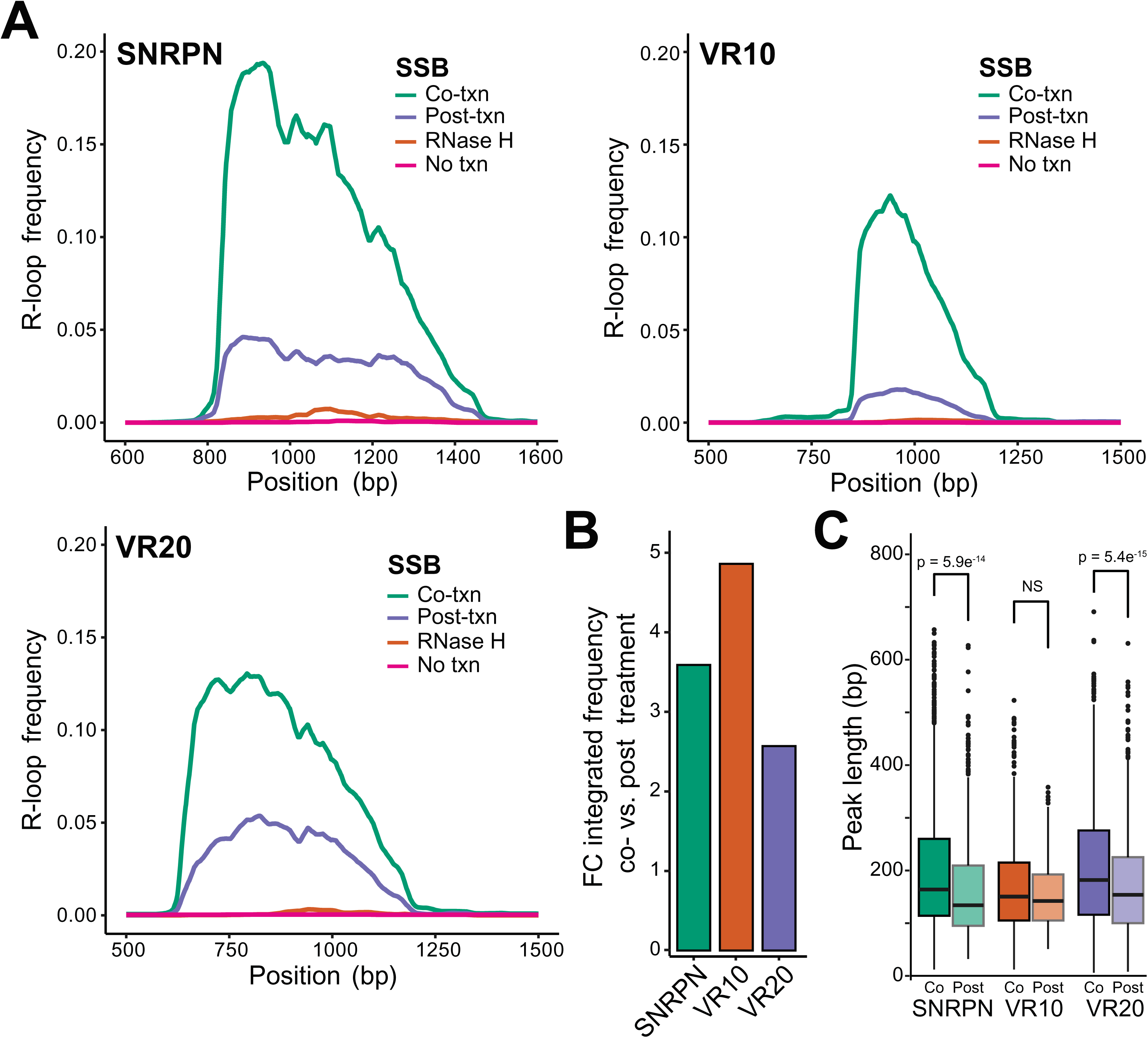
**A:** R-loop frequency plots calculated from SMRF-seq data for linearized SNRPN, Airn, VR10 and VR20 containing plasmids. **B:** Boxplot of R-loop peak length measured via SMRF-seq comparing co and post transcriptional SSB treatments for the four previously mentioned linearized plasmid substrates. P values calculated using Student’s t-test. **C:** Boxplot of fold change in R-loop frequency by base pair between co and post SSB treatment of the VR containing region from the VR-10 and VR-20 plasmids.

We measured the lengths of R-loops formed under co- and post-transcriptional conditions to determine whether SSB also allows the growth of R-loops during R-loop elongation. Median R-loop lengths under post-transcriptional SSB addition ranged from approximately 120 to 150 bp, consistent with prior findings (12). A modest increase in length was observed for two out of the three tested constructs (**Fig 1C**). This indicates that stabilization of the non-template strand may allow for slightly more R-loop extension compared to an un-stabilized non-template strand. The effect size, however, was small compared to the increase in overall R-loop frequency.

### Transcription through nicks increases R-loop formation regardless of sequence or distance from promoter

Nicks may allow the displaced DNA strand to locally dissociate from the DNA template strand, thereby favoring the reannealing of the nascent RNA to initiate an R-loop (20). Alternatively, or in addition, we propose that R-loops that initiate at, or in the immediate vicinity of nicks would not have to “pay” the energetic cost imposed by the formation of a Y junction. An R-loop initiated downstream of a nick indeed does not have to contend with a start junction due to the interruption of the phosphate backbone. If correct, this hypothesis predicts that R-loop formation following a nick should be less constrained by the favorability of the downstream DNA sequence, due to the lower requirement for energy return from RNA:DNA hybrid formation.

To test this hypothesis, we utilized a CRISPR-Cas9 nickase system (21), (22) and three single guide RNAs (sgRNAs) (23) to introduce DNA nicks in the non-template strand of an R-loop-prone plasmid carrying the human *SNRPN* gene (12, 24). These sgRNA molecules, termed sgRNA 1, 2, and 3 directed strand-specific Cas9-mediated nicks 91, 569, and 973 bp downstream of the T7 transcription start site (TSS), respectively (**Fig 2A**). Analysis of the local average R-loop energy for the first 100 bp downstream of each nick, showed that the sequence following sgRNA 2 was by far the most favorable (**Fig S2A**), corresponding to a well-known GC-skewed R-loop formation hotspot (12). The region following sgRNA 1 was moderately favorable, while the region downstream of sgRNA3 was the least favorable and the furthest downstream from the promoter.

**Figure 2.**
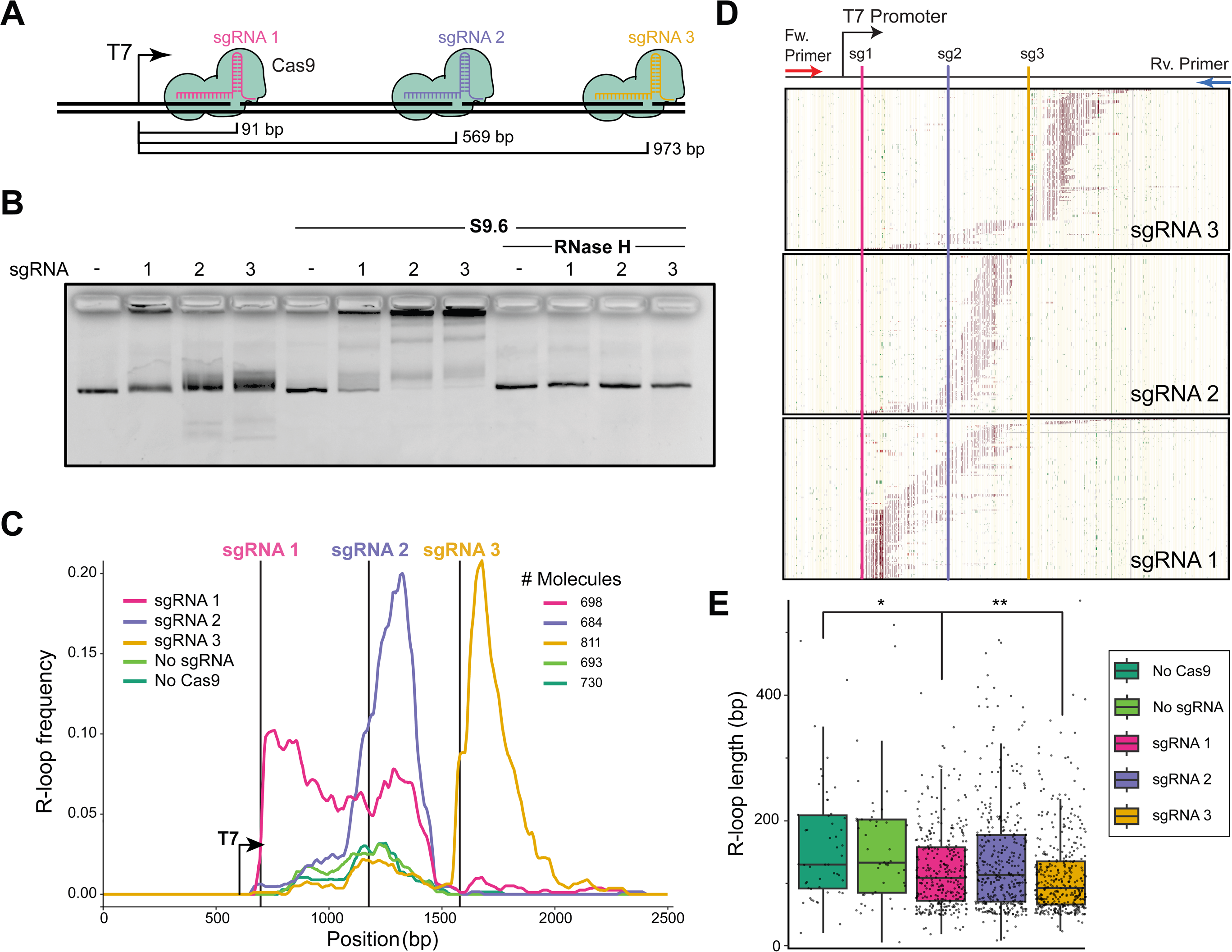
**A**: Diagram showing target sites of the three sgRNAs used to induce Cas9 strand-specific DNA nicks relative to the T7 transcription start site. **B**: Agarose gel electrophoresis of *in vitro* transcription reaction on nicked and unnicked linearized plasmids. **C**: Frequency plot of SMRF-seq signal from *in vitro* transcription reactions on the non-template DNA strand. Locations of Cas9 nicks are shown as vertical lines; the position of the T7 promoter is indicated. The number of independent molecules analyzed by SMRF-seq is indicated. **D:** A representative sampling of SMRF-seq footprints collected for each nicked plasmid is displayed. 200 independent R-loop-carrying molecules are displayed vertically stacked. The position of cytosines is shown by yellow vertical bars. Red tick marks indicate cytosines that were single-stranded and part of an R-loop footprint. Green tick marks indicate cytosines that were converted but not considered to be part of a footprint. **E**: Boxplot of R-loop lengths measured in base pairs for nicked and unnicked conditions.

The induction of Cas9-mediated nicks in supercoiled plasmid templates was observed via a characteristic loss of mobility during agarose gel electrophoresis (**Fig S2B**). Only reactions showing at least 95% efficiency were utilized in downstream experiments. The location of the nicks was verified using Sanger sequencing by a tell-tale early termination of the sequencing reaction at the expected nick site (**Fig S2C**). Having validated a robust Cas9-based nick induction methodology, we next qualitatively determined the impact of nicks on R-loop formation in bulk by performing *in vitro* transcription reactions utilizing T7 RNA polymerase followed by electrophoretic mobility shift assays. All substrates were deproteinized and linearized prior to transcription to remove any effect of DNA topology. R-loop formation was inferred by a characteristic upward gel shifting. Treatment with sgRNA 1, 2 and 3 resulted in increased upward shifting that was greatly accentuated by incubation with the anti-RNA:DNA hybrid S9.6 antibody (25) prior to electrophoresis (**Fig 2B**). We confirmed that gel shifting reflected R-loop-mediated phenomena by treating transcribed substrates with purified RNase H prior to gel electrophoresis. This treatment completely eliminated S9.6 binding. Thus, transcription of nicked substrates causes a strong increase in R-loop formation, extending prior results (20). Interestingly, all three nicked substrates responded strongly to nicks when transcribed, suggesting that the distance from the TSS to the nick, and the local DNA sequence, do not qualitatively influence the impact of a nick on R-loop formation.

### SMRF-seq reveals R-loop induction at or in the immediate vicinity of nicks

To provide quantitative and spatial insights into the effect of non-template strand nicks on R-loop formation, we subjected *in vitro* transcription reaction products to a modified SMRF-seq protocol which includes a post-bisulfite treatment nick repair step to allow for successful PCR amplification of nicked templates (**Fig S2D**; see SI Appendix). Importantly, SMRF-seq does not involve any enrichment for R-loop-carrying molecules and instead samples all plasmids in the reaction, providing an unbiased, quantitative, high-resolution, and strand-specific measurement of R-loop formation on single DNA molecules at ultra-deep coverage.

Transcription through unnicked, linear templates resulted in a small number of R-loops that tended to cluster over the most favorable, GC-skewed, portion of the transcribed region, in agreement with previous observations (12). This R-loop prone region maps to the area targeted by sgRNA 2. *In vitro* transcription after treatment with Cas9 alone in the absence of sgRNA did not result in any significant change in R-loop frequency, as expected (**Fig 2C**). In stark contrast, the bulk R-loop frequency was dramatically increased immediately at, and downstream of, the site of Cas9-induced nicks (**Fig 2C**). For sgRNA 2 and 3, a sharp peak of R-loop formation was observed over a ∼400 bp region downstream of the nicks. For sgRNA 1, increased R-loops were induced over a wider region ∼800 bp downstream of the nick. RNase H treatment nearly eliminated all SMRF-seq signal confirming that it originated from *bona fide* R-loops (**Fig S2E**). Additionally, the frequency increases observed here were highly strand-specific, with no effect observed on the template DNA strand (**Fig S2F**), as expected from R-loops.

Analysis of individual DNA molecules recovered from SMRF-seq confirmed the presence of clear R-loop footprints downstream of the nick sites (**Fig 2D**). We determined that the structures formed downstream of the sgRNA target sites resulted from transcription events originating at the T7 promoter, since adventitious loading of the T7 RNAP at nicks did not result in significant transcription or R-loops (**Fig S2G**). R-loop lengths were measured from individual R-loop start and stops. For unnicked samples or Cas9-only treated samples, median R-loop lengths were 173 bp. Nick-induced R-loops were slightly smaller (**Fig 2E**), with the most significant difference observed for R-loops induced by sgRNA 3, and to a smaller extent, sgRNA 1. As mentioned, DNA sequence energetics for the sgRNA 3 nick-downstream region was the least favorable for R-loop formation, possibly accounting for the reduced R-loop size.

### DNA nicks stimulate R-loop formation by one to two orders of magnitude, regardless of nicking methodology

We measured R-loop frequencies over a 50 bp window immediately downstream of each nick in both nicked samples and unnicked controls. R-loop frequency increased a stunning 155-fold for the sgRNA 3 site, while sgRNA 1 and 2 led to 22-fold and 6-fold increases, respectively (**Fig 3A**). The main reason for this disparity in fold-change was due to the differences in background R-loop favorability in unnicked samples. R-loop frequencies in nicked samples were relatively consistent, ranging from 9 to 16% between sgRNAs (**Fig 3A**). This suggests that R-loop induction at nicks is similar regardless of the distance to the TSS, contrary to a previous report (20). This implies that the length of the trailing RNA transcript when the polymerase reaches the nick does not have an appreciable effect on the nick-induced R-loop frequency. It also implies that R-loop induction following a nick can be extraordinarily effective, even when the downstream DNA sequence is not intrinsically favorable to R-loops, as observed for sgRNA 3.

**Figure 3.**
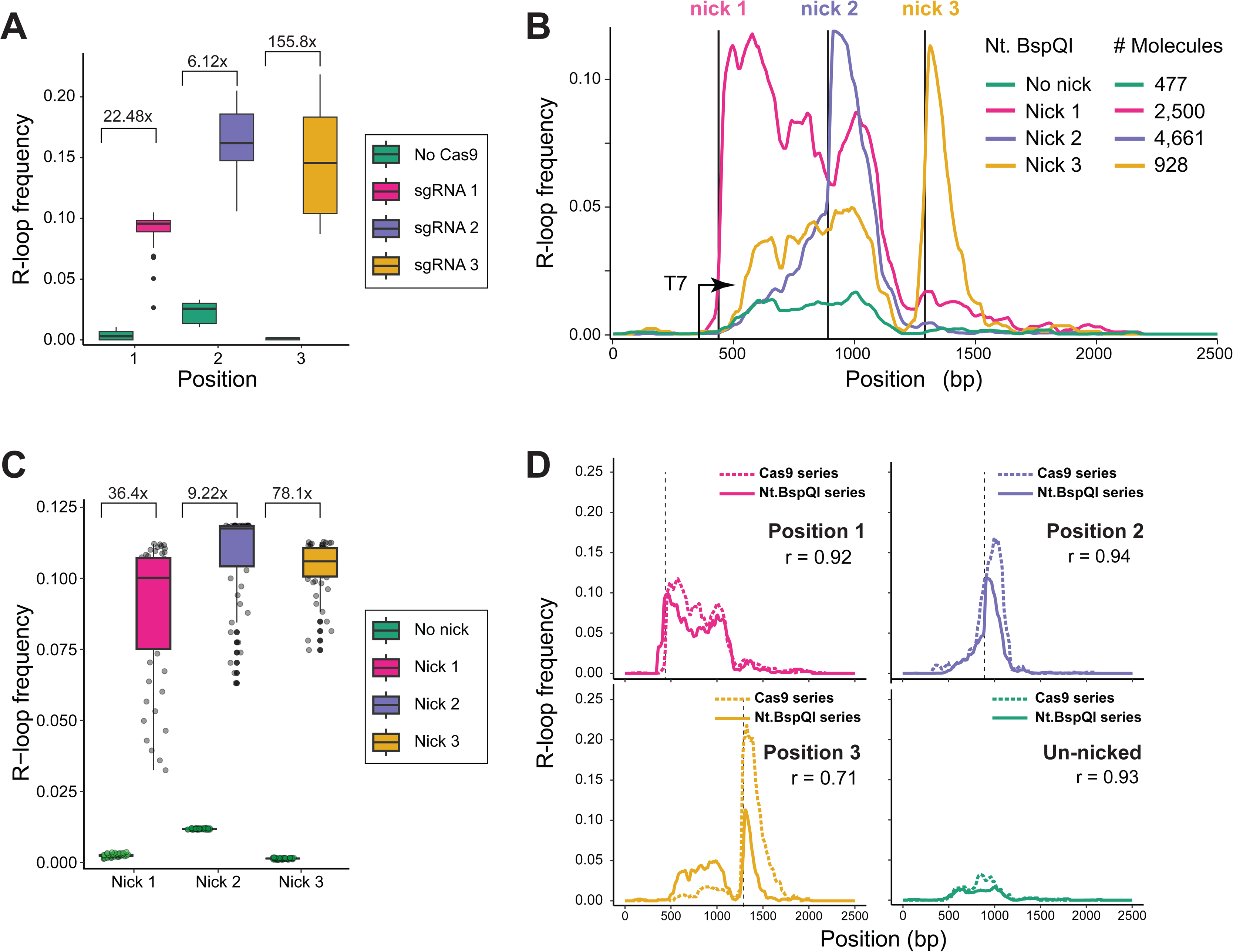
**A**: Box plot of R-loop frequencies from SMRF-seq measured over the 50 bp region immediately following the site of Cas9-induced nicks in nicked vs. unnicked samples. The fold-change between unnicked and nicked samples is indicated above paired samples. **B**: R-loop frequency plot from SMRF-seq data for linearized plasmids with or without Nt.BspQI-induced nicks. Horizontal lines indicate the location of Nt.BspQI recognition sites. The number of independent molecules sampled is indicated at right. **C**: As in panel A, except data is for Nt.BspQI-induced nicked and unnicked samples. **D:** Comparison between R-loop frequency plots calculated from SMRF-seq signal between analogous Nt.BspQI and Cas9 nicked and unnicked samples. The plots indicate the Pearson correlation coefficient between the two curves.

To verify that the increase in R-loop frequency at nicks was not Cas9-specific, we engineered the *SNRPN* region to contain an Nt.BspQI recognition site either 126, 578, or 978 bp downstream of the T7 promoter. Nt.BspQI treatment allowed us to generate site-specific nicks at high efficiency in positions immediately adjacent to that of the Cas9-induced nicks. Nt.BspQI-induced nicks greatly increased R-loop formation compared to unnicked controls in a manner highly similar to that observed for Cas9-induced nicks (**Fig 3B**). The R-loop frequencies downstream of each nick site were similar for all three positions and the fold changes ranged from 9-fold for Nick 2, to 36-fold for Nick 1 and 78-fold for Nick 3 (**Fig 3C**). As observed previously, nick-induced R-loops were highly specific to the non-template strand (**Fig S3A**), were not apparent in nicked but un-transcribed samples (**Fig S3B**) and were abrogated by RNase H treatment (**Fig S3C**). Direct comparison of R-loop frequency curves confirmed a high correspondence between Nt.BspQI-induced and Cas9 nickase-induced samples (**Fig 3D**), establishing that nicks are powerful R-loop initiators regardless of nicking strategy or distance from the TSS.

The ability of nicks to stimulate R-loop formation regardless of the downstream sequence is consistent with the idea that the energetic cost of sustaining a Y-junction is the major bottleneck to structure formation (12). Our new dataset provides an opportunity to adapt our R-loop equilibrium energy model, R-looper, to simulate nick-induced R-loop formation and to empirically estimate the junction energy. We found that the current values, based on a simple strand separation model (26), likely under-estimate the R-loop junction energy. The best fit between predicted and experimental data was observed at a junctional energy (a) value of 17.9 kcal/mol, representing a 1.79x fold increase compared to the previously utilized (a) value (see SI appendix). Importantly, the modeling confirmed that simply relieving the junction energy at one base-pair can result in a dramatic shifting of R-loop distribution to that position, in effect creating a major R-loop hotspot (see SI appendix).

### Direct visualization of R-loops formed at nicks reveal unique and complex “forked” secondary structures

Atomic force microscopy (AFM) of *in vitro* generated R-loops has been used previously to characterize the resulting structures (27). Given the significant increases in R-loop frequency observed at nicks, we wanted to determine if these structures resembled those formed in the absence of nicks. A total of 287 molecules were imaged, including 144 Nt.BspQI-nicked and 143 unnicked molecules. The median contour length of all molecules was estimated to be 3473 bp (1181 nm), a 2.3% deviation from the expected plasmid length (**Fig S4A**). As expected, untranscribed molecules, whether nicked or unnicked, did not carry any structural feature deviating from that expected from B-DNA. This confirms that nicks in the absence of transcription did not induce secondary structures. Structural features deviating from B-DNA were observed on approximately 30% of transcribed molecules (**Fig 4A**). Within this subset, three distinct classes of features were observed and categorized as loops, blobs, and forks (**Fig 4B** and **Fig S4B**). Only one feature was detected per molecule. Loops and blobs have previously been reported as *bona fide* R-loop features (27); “forks”, however, are novel transcription-induced features. Forks are characterized by the appearance of an unexpected segment of double-stranded nucleic acid jutting from the main molecule and seemingly connected by what often appears to be ssDNA (**Fig 4B**). Some fork objects also displayed various degree of complexity, with the appearance of branched structures (**Fig S4B**).

**Figure 4.**
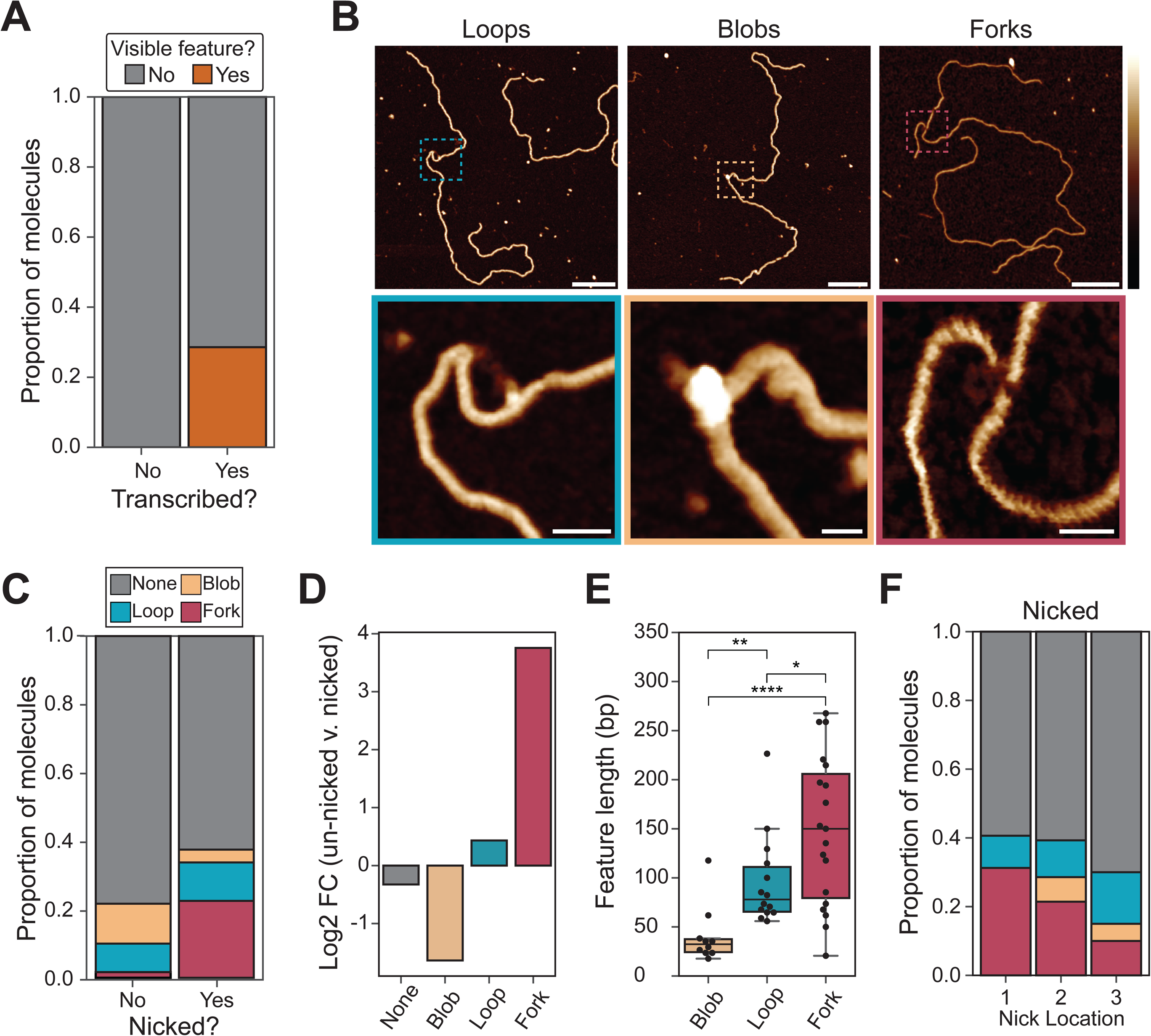
**A:** Barplot showing the proportion of molecules with any visible feature type observed using AFM on transcribed and un-transcribed molecules. **B:** Representative AFM images of three observed structures classes; forks, loops and blobs. Scale bars on upper images represent 100 nm. Lower image scale bars are equal to 10 nm. Height scale = -1 to 3 nm. **C:** Barplot showing the proportion of each feature type (or lack of feature) observed via AFM on transcribed nicked and unnicked samples. **D:** Barplot showing fold change between proportion of each feature type between unnicked and nicked transcribed molecules observed using AFM. **E:** Boxplot showing distribution of feature lengths by type in base pairs. **F:** Boxplot showing proportion of feature types observed for each nick location.

Transcription through unnicked plasmids produced a large number of loop and blob features, consistent with prior observations (**Fig 4C**) (27). 21.6% (13 molecules out of 60) of all transcribed unnicked molecules carried a visible R-loop feature; only one molecule carried what appeared to be a “fork”. Transcription through nicked samples resulted in a significant increase in the overall proportion of molecules carrying a feature: 33.8% (27 molecules out of 80) of transcribed nicked molecules carried a visible non-B DNA feature. Thus, nicks induced a 1.74-fold increase in the frequency of non-B DNA features detected by AFM. In sharp contrast to unnicked samples, the majority of molecules resulting from transcription through nicked substrates carried fork features (**Fig 4C,D**). Overall, there was a dramatic ∼13-fold increase in the number of fork features between nicked and unnicked transcribed samples, while the proportion of blobs decreased nearly 3-fold (**Fig 4D**). Features in nicked samples were significantly longer than those observed in unnicked samples; this increase was primary driven by the fact that fork objects are larger than blobs or loops (**Fig 4E**). Forks had lengths averaging 149 bp (+/- 76 bp), while loops and blobs had average lengths of 96 bp (+/- 46 bp) and 41 bp (+/- 30 bp), respectively. The relative proportion of fork objects varied depending on the nick position, with nick position 1 showing the most forks, and nick position 3 the least (**Fig 4F**). Overall, fork objects represent a novel nick-induced, transcription-dependent, class of R-loop structures.

### Fork structures most likely result from non-template strand “peeling” and self-folding during R-loop formation

We hypothesized that the unique and unexpected fork objects could be explained by the non-template strand peeling from the template downstream of the nick, as it is not covalently attached to the upstream-of-nick non-template strand. We further hypothesized that this strand may fold back onto itself forming a variety of possible hairpin-like structures that would result in a dsDNA character when observed via AFM (**Fig 5A**). Further examples of fork structures with corresponding inferred strand positions and orientations are shown in **Fig 5B**. To test this hypothesis in bulk, we transcribed nicked substrates and treated them simultaneously by RNase H and low levels of nuclease S1. We reasoned that if the RNA portion of the RNA:DNA hybrid was rapidly removed by RNase H, the fork structure would likely remain intact for some time due to the additional stability afforded by its secondary structure. RNase H digestion would therefore leave a region of exposed ssDNA on the template strand with the approximate size of the RNA:DNA hybrid (**Fig S5A**). This ssDNA should be hypersensitive to S1 digestion. The self-paired fork structure itself should be sensitive to S1 cleavage in places where ssDNA is exposed. Two products are expected. One corresponds to a left fragment of defined size running from the end of the linear molecule to the nick. The second right fragment should be of variable size depending on the position and length of the RNA:DNA hybrid and the geometry of the self-paired fork (**Fig S5A**). These digestion products should be apparent by agarose gel electrophoresis given a high enough concentration of forks were present. Regular R-loops formed outside of a nick would be expected to resolve back to dsDNA immediately after the RNA strand is digested by RNase H.

**Figure 5.**
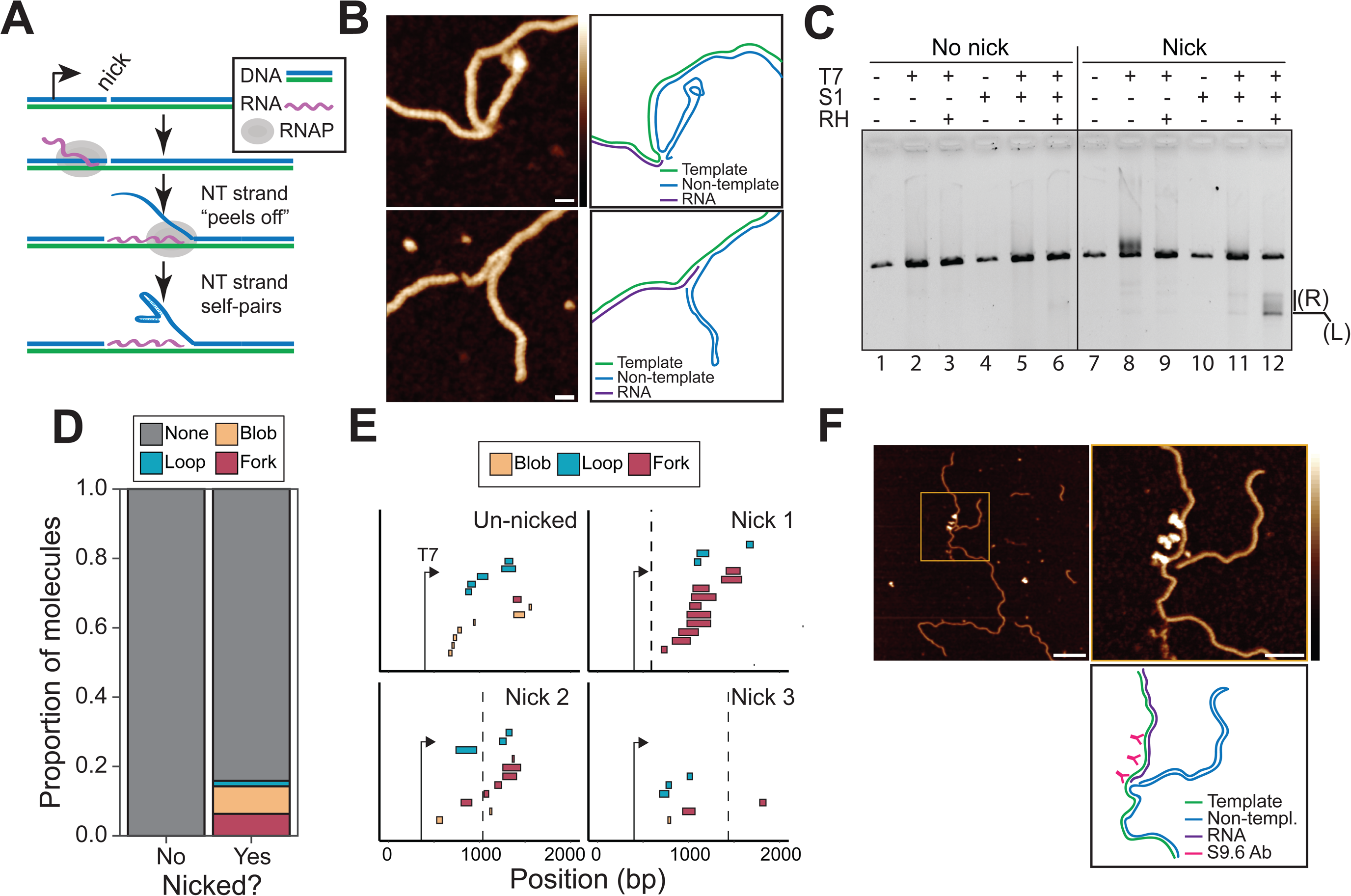
**A:** Cartoon diagram of model of an R-loop associated fork structure at the site of a non-template strand nick. **B:** AFM images and corresponding strand tracing diagram models of two fork structures observed on transcribed and nicked substrates. Scale bar = 10 nm. Height scale = -1 to 3 nm. **C:** Agarose gel electrophoresis of an *in vitro* transcription reaction on linearized nicked and unnicked SNRPN containing plasmid. Pluses or minuses above lanes indicate the addition or absence of a particular enzyme in the reaction. The symbols T7, S1 and RH correspond to T7 RNA polymerase, S1 nuclease and RNase H1 (E. Coli) respectively. **D:** Barplot showing the proportion of feature types observed via AFM in nicked and unnicked transcribed and Rnase H1 treated molecules. **E:** Footprint plot showing the locations of features with respect to each nick location and the T7 promoter seq. The expected nick site is indicated by a vertical dashed line. **F:** AFM image and strand tracing cartoon model of a nicked and transcribed plasmid treated with S9.6 antibody. Scale bar on left image = 100 nm. Scale bar on right image = 40 nm. Height scale = -1 to 3 nm.

As expected, we found minimal S1 nuclease digestion products in the unnicked transcribed and RNase H-treated sample (**Fig 5C**, lane 6), consistent with rapid R-loop resolution back to dsDNA upon RNase H treatment. Transcription of the Nt.BspQI-nicked substrate led to significant upward smearing compared to an unnicked substrate (compare lanes 2 and 8), consistent with increased R-loop formation. S1 treatment of transcribed nicked molecules led to the disappearance of the smear (compare lanes 11 and 8), consistent with it being due to secondary structure formation on the displaced strand, as proposed (27). Co-treatment with S1 nuclease and RNase H led to the clear appearance of new digestion products compared to S1 treatment alone (compare lanes 12 and 11), indicating that the new ssDNA substrates for S1 nuclease were generated by RNase H activity, as hypothesized. The products appeared as a lower band and an upper smear (**Fig 5C**). The lower product closely corresponded in size to the distance expected between the BsaI site used to linearize the plasmids and the Nt.BspQI target site. The defined character and size of this band are entirely consistent with this product corresponding to the expected left product (**Fig S5A**). The upper smear, in turn, is consistent with the products expected from S1 nuclease cleavage of forks of variable structures and of RNase H-exposed ssDNA of variable position and lengths originating from the right side of plasmid substrates. There was minimal visible S1 digestion on nicked untranscribed samples (lane 10), indicating that the digestion products were both transcription- and RNase H-dependent, as expected.

Our hypothesis for the origin of fork objects makes three further predictions. First, RNase H treatment of fork objects may not result in a full return to dsDNA due to the intrinsic stability of the self-paired non-template strand. Second, fork objects should appear after nicks. Third, the ability of a given region to form fork objects should depend on the ability of the region to undergo self-folding. To test the first prediction, we applied AFM on RNase H-treated nicked and unnicked transcribed plasmids. As expected, no visible structures were observed in any of the RNase H-treated unnicked samples (61 total molecules), consistent with complete resolution to B-form dsDNA (**Fig 5D**). However, a relatively large proportion of residual structures persisted in the nicked samples post RNase H treatment. These residual structures were mostly composed of blobs and forks (**Fig 5D**). The presence of fork objects post RNase H treatment is consistent with the notion that self-paired structures on the non-template strand may remain stable and resist unwinding and reannealing to the template strand. It is possible that blob objects represent collapsed forks, although this hypothesis requires additional investigation. To test the second prediction, we plotted the location of all observed structures for the three nicked and unnicked conditions after AFM imaging and careful recording of the position of each structure relative to the ends of the molecule. We found that, in general, forks tended to be located post nicks (**Fig 5E**). For nick 1, all structures were found post nick although we note that the distance from the T7 promoter to the nick is short in this configuration. For nick 2, 5 out of 6 fork objects were located post nick. For nick 3, only one fork object was observed, and it was located prior to the nick. Quantification of this data showed that forks showed a clear bias in their position, occurring in 80% of the cases after the nick position. By contrast, blobs and loops showed no biases in their location and were near equally likely to occur before or after nick sites (**Fig S5B**). From this, we conclude that fork objects typically occur downstream of nick sites. To test the third prediction, we noted that the proportion of fork objects was highest for Nick 1 compared to Nick 2, and lowest for Nick 3 (**Fig 4F**). We used secondary structure prediction algorithms to determine the self-folding probability of regions downstream of each nick and observed the sequence around Nick 1 had the lowest (most favorable) predicted folding energy while Nick 2 showed an intermediate prediction and Nick 3 is positioned in a relatively unfavorable region for folding (**Fig S5C**). Overall, these results are consistent with our hypothesis that fork structures arise by peeling and self-folding of the non-template strand during R-loop formation.

As a final test of the hypothesis, we wished to test for the presence of an RNA:DNA hybrid portion next to the fork object itself. Since AFM alone can’t easily differentiate RNA:DNA hybrid from dsDNA, we took advantage of the S9.6 anti-RNA:DNA hybrid antibody to mark hybrids and performed AFM imaging on nicked and unnicked substrates. As expected, we observed minimal S9.6 binding to plasmid molecules in the untranscribed nicked condition, indicating absence of RNA:DNA hybrids or R-loops. Clear binding was observed on unnicked transcribed plasmids (**Fig S5D**). For nicked substrates, we were able to observe clear instances where S9.6 binding was located immediately adjacent to the protruding fork, but was absent from the fork itself (**Fig 5F**), as predicted if the fork feature itself was a dsDNA structure immediately adjacent to an RNA:DNA hybrid.

## DISCUSSION

R-loop formation is influenced by the favorability of the underlying DNA sequence and by the level of negative DNA superhelicity available (12, 28). Our work here reveals that the non-template DNA strand exerts profound additional impacts on R-loop formation independently of DNA sequence or topology. Consistent with the notion that the nascent RNA competes with the non-template DNA strand for binding to the template DNA strand during R-loop formation (20), we show that binding of the non-template DNA strand by *E. coli* SSB strongly stimulated R-loop frequency by up to 5-fold. Importantly, this effect required addition of SSB co-transcriptionally such that SSB could interact with the displaced strand of R-loops as soon as it became available. Furthermore, SSB was not necessary to ensure R-loop stability after the transcription reaction was complete. We interpret this to mean that SSB stabilized short nascent R-loops that would have otherwise collapsed back to duplex DNA during early R-loop formation. This is consistent with the view that forming an R-loop involves a significant energetic cost in the form of creating at least one Y-junction at the point where the nascent RNA invades the duplex DNA. This energetic cost can be “repaid” by the formation of a more stable RNA:DNA hybrid and by the relaxation of negative supercoiling (12). However, this energetic return from RNA:DNA hybrid formation and superhelical relaxation scales with size, rendering short R-loops particularly vulnerable to early collapse. SSB can bind to as little as 35 nucleotides of ssDNA per tetramer (29), ensuring that binding to the displaced non-template strand can occur early. SSB binding is expected to increase as more ssDNA becomes available during R-loop elongation, resulting in the stabilization of the displaced DNA strand and in a reduced tendency for R-loop collapse. According to this model, the effect of SSB should only be manifested in the stabilization of R-loops that have already initiated and for which an ssDNA platform is available. Indeed, while SSB significantly increased R-loop frequency, it did not shift the locations of R-loops. SSB binding only had a minor effect of R-loop lengths suggesting that its main impact is on R-loop initiation. Overall, this work shows that R-loop formation is highly dynamic and that in the absence of stabilizing factors, most R-loops that initiate end up in spontaneous early collapse due to the reannealing of the non-template DNA strand. Our findings suggest that single-stranded DNA binding proteins may play an important role in regulating R-loop formation in cells. Conditions where SSB or RPA availability is limiting, including conditions of replication stress, genotoxic damage (30, 31), or unprocessed RNA accumulation (32), might be expected to show reduced R-loop loads due to early collapse. Likewise, it is possible that RPA availability may regulate the resolution of R-loops by Ribonuclease H1 given the proposed role of RPA in recruiting this R-loop resolvase (18).

Our work further demonstrates that nicks in the non-template DNA strand can increase R-loop frequencies by up to two orders of magnitude, dramatically extending previous observations (20).Our findings further demonstrate that the effect of nicks is largely independent of both the underlying DNA sequence and the distance of the nick to the transcription start site. The impact of nicks on R-loop frequencies can be most easily explained by the high energetic cost of creating a Y-junction when initiating an R-loop. If an R-loop instead initiates at a nick, the cost of this junction is cancelled. Indeed, our data clearly show that the source of increased R-loops in nicked templates arises from R-loop initiation at, or in the immediate vicinity, of the nick (**Figs 2, 3**). Thus, unlike the addition of SSB-type proteins that increase R-loop frequencies without affecting the location of R-loops, nicks profoundly change the R-loop landscape and can transform a nearly inactive sequence into a major R-loop hotspot. Our proposal that this effect is due to the cancellation of the junction energy for nick-initiated R-loops is supported by mathematical modeling. For junction energy costs around 18 kcal/mol, the modeling data clearly shows that predicted R-loop locations dramatically shift from the most favorable regions based on DNA sequence, to the position of the nick itself (**SI Appendix**).

The potential for non-template strand DNA nicks to act as strong initiators or R-loop formation *in vitro* is intriguing in the context of diseases associated with increased DNA nicks. Aicardi-Goutières syndrome (AGS), for instance, is caused by inherited mutations in nucleic acid metabolism genes (33–35) including the genes encoding for all three subunits of ribonuclease H2. RNase H2 is the primary enzyme responsible for ribonucleotide excision repair (RER), a critical pathway that removes single ribonucleotides (rNMPs) embedded in genomic DNA (36, 37). Loss of RNase H2 and AGS mutations result in accumulation of rNMPs in yeast, murine, and human genomic DNA and cause a secondary increase in DNA nicks due to the aberrant activity of DNA topoisomerase 1 at rNMPs (38). Interestingly, genome wide mapping of RNA:DNA hybrids in AGS patient-derived fibroblasts revealed significant increases in RNA:DNA hybrid levels. Such increases, however, were relatively unique to each patient sample and were observed over DNA sequences typically devoid of favorable R-loop-forming characteristics (39). It is plausible that the sporadic accumulation of DNA nicks in RNase H2-mutated cells may cause nick-induced RNA:DNA hybrid formation. Mutations in the *ATM* gene cause Ataxia-Telangiectasia (40) and were also shown to cause increased single-strand break loads due to elevated reactive oxygen species (ROS) (41, 42). Intriguingly, ATM mutant cells also exhibit higher R-loop levels and this increased R-loop burden can be suppressed by antioxidant treatment (41) , consistent with a role for ROS-induced nicks in promoting RNA:DNA hybrid accumulation. Further observations are required to determine if the effect of DNA nicks on R-loop formation described here *in vitro* applies in human cells and becomes exacerbated in disease conditions. Two observations support that nicks may indeed serve as initiation sites for R-loops in cells. First, treatment of mammalian cells with the DNA topoisomerase I poison camptothecin results in a rapid increase in DNA nicks and a burst of R-loops (43, 44). Second, recent work established that strand discontinuities such as unprocessed flaps or un-ligated Okazaki fragments can result in a secondary accumulation of DNA:RNA hybrids in yeast (45).

In addition to increasing the local frequency of R-loop formation, nicks also appear to cause the formation of a unique type of R-loop “fork” structures. Direct visualization of R-loops by AFM revealed that nicks cause a 13-fold increase in such fork structures. Structure mapping, enzymatic probing, and antibody binding experiments support the notion that these structures arise from peeling and folding back of the non-template DNA strand as a result of the formation of an RNA:DNA hybrid. In addition, our data supports that regions more prone to self-folding showed more fork structures, while regions least prone to it showed the least structures. The resolution of nick-induced forked R-loops is expected to be much more demanding than that of normal R-loops. In the latter case, RNase H activity or an RNA:DNA helicase would suffice to restore B DNA once the RNA:DNA hybrid has been removed. Forked R-loops, however, would require additional factors to resolve the fork structure, including DNA helicases to unwind self-paired hairpins, an annealing activity to pair the previously peeled off non-template strand back to the template DNA strand, and a DNA ligase to seal the nick. The faithful resolution of forked R-loops will also require a precise choreography of enzymatic activities. If, for instance, the RNA:DNA hybrid is resolved prior to the unfolding and reattachment of the fork, a long ssDNA gap will be left on the template strand. Similarly, improper cleavage of the forked structure by a flap endonuclease may lead to the formation of a two-stranded RNA:DNA hybrid, and likely to a ssDNA gap. It is therefore likely that nick-induced forked R-loops may represent a particularly genotoxic class of R-loops that threaten genome integrity.

## MATERIAL AND METHODS

For more information on plasmid substrates, *in vitro* transcription assays, nicking procedures, single-molecule R-loop footprinting, AFM imaging, and data analysis, please consult the SI Appendix.

## ACKNOWLEDGMENTS.

We thank members of the Chedin laboratory for critical feedback and valuable discussion. This work was supported by National Institutes of Health grant R35 GM139549 and National Science Foundation grant NSF/DMS 2054347 to F.C. We wish to acknowledge a UKRI Future Leaders Fellowship MR/W00738X/1 to A.L.B.P, and the Henry Royce Institute for Advanced Materials EP/R00661X/1, EP/S019367/1, EP/P02470X/1 and EP/P025285/1 and Xinyue Chen for Dimension FastScan access and support through Royce@Sheffield. We thank Dr. S.C. Kowalczykowski for the kind gift of purified *E. coli* SSB protein.

## Supporting Information Appendix for

### Supporting Information text

#### Estimation of the energetic cost of the R-loop junctional energy

Previously, R-loop modeling using R-looper (1) assumed that the junction energy parameter (*a*) was 10 kcal/mol, following earlier experimental measurements for B/Z transitions and local strand separation (2–4). The junction energy for R-loops has not been experimentally determined but could be higher than simple strand separation given that R-loops must accommodate the RNA strand of the RNA:DNA hybrid inside a strand-separated bubble. To empirically model (*a*), we modified R-looper such that no junction energy would be applied specifically for R-loops initiating at nicks. We assumed that negative superhelicity was -7% for these calculations, consistent with measurement of transcription-induced dynamic supercoiling (5). The junction energy itself was varied systematically and R-looper was used to generate simulated R-loop footprints. Focusing on the distal position 3 for which a nick caused a two-order of magnitude increase in R-loop frequencies, we observed that for a default (*a*) value of 10 kcal/mol, most R-loops are predicted to form around the central, energetically favorable region, regardless of whether junction energy relief was applied at the nick (**Fig S6A**). As the value of (a) is increased, however, the cost of initiating R-loops there is raised. By contrast, initiation at the nick, where the increasing junction energy cost does not apply, is becoming progressively more likely, outweighing the penalty associated with initiating in a poor R-loop forming DNA sequence. At an (a) value of 17.9 kcal/mol, we observed the best fit for simulated vs. observed R-loop footprints (**Fig S6B**), as measured by the Precision value. Under that condition, most R-loops are now predicted to form at the nick, simulating a dramatic shift of R-loop positions. This data therefore suggests that the critical junction energy value (a) is significantly higher than previously thought. We note, however, that the (a) value estimate is likely to vary with the local DNA sequence and changes considerably with the level of available negative DNA superhelicity. As the negative supercoiling is increased, the value of (a) needed to shift the distribution of R-loops to the nick sites is also strongly increased (**Fig S6C**). This is consistent with the notion that to observe shifting to the nick, one must increasingly penalize the formation of R-loops over the most favorable regions based on DNA sequence and topology. Regardless, our data and modeling suggest that the value of (a) has been previously under-estimated and that nicks derive their ability to focus R-loop initiation from the relief of this otherwise severely limiting parameter.

### Materials and methods

#### Plasmid substrates

The VR-10 and VR-20 plasmids were cloned by digesting the pFC9 SNRPN containing plasmid (1, 6) with SacI-HF® (NEB, R3156S) and EcoRI-HF® (NEB, R3101S) according to the manufacturer’s online protocol. The large fragment resulting from the digest was then separated via agarose gel electrophoresis and subsequently extracted. Synthetic DNA sequences, or variable regions (VRs) with corresponding pFC9 homology sites of 250 bp in length were then inserted via Gibson assembly (7) into the pFC9 based vector. The VR sequences were designed to have specific R-loop forming properties, namely controlled GC and content. VR-10 was designed to have an overall GC skew and content of 0.2 and 0.5 while VR-20 was designed with 0.4 and 0.6 respectively. All guanines within the VR-20 sequence were clustered into triplets and distributed at a regular interval through the sequence while the guanines in VR-10 were allowed to distribute randomly. The VR-10 and 20 plasmid sequences were confirmed via Sanger sequencing.

#### Plasmid isolation

Plasmids were transformed into *E. coli* DH5α chemically competent cells and single colonies were grown on LB ampicillin agar plates. Single colonies were selected and grown in 300 ml LB culture with 100 µg / ml of ampicillin in 1 L flasks at either 30° or 37°C overnight with shaking at 200-250 rpm. Negatively supercoiled plasmid substrates were purified by alkaline lysis protocol followed by column cleanup (Qiagen) and eluted into either TE or EB buffer.

#### *In vitro* transcription reactions

*In vitro* transcription assays were performed as described in Ginno *et al.*, (2012) (6) with 20 ng / μl DNA and either T7 (NEB M0251S) or T3 (NEB M0378S) RNA polymerases (2.5 U / μl) in the buffer provided by the polymerase manufacturer supplemented with rNTP mix solution (NEB, N0466S) added to a final concentration of 0.5 mM and DTT (NEB, B1222A) to a final concentration of 10 mM. Reactions were incubated at 37°C for 20 minutes and then were terminated by the addition of EDTA to a final concentration of 35 mM or by physical separation of the promoter from the transcribed region by restriction digest. For direct visualization, the reaction products were treated with RNase A (33 ng/ml) for 30-45 minutes at 37°C to degrade excess free RNA. To confirm presence of R-loops, a portion of the transcribed sample was treated with RNase H (NEB M0297S, 40 ng DNA/1U) for 30 minutes at 37° C to degrade RNA:DNA hybrids. The products were then separated by agarose gel electrophoresis using 0.8% 1x TBE gel at a constant 60 volts for 2 hours. The gels were then post stained with ethidium bromide and imaged with a BioRad GelDoc EZ imager.

#### Design and synthesis of single guide RNAs

Specific Cas9 single guide RNAs (sgRNAs) target sites were selected using Benchling online software. Oligos with the selected target sites were ordered according to the NEB EnGen® sgRNA Synthesis Kit, *S. pyogenes* (E3322V) kit protocol. sgRNAs were then synthesized according to NEB protocols and purified via phenol-chloroform ethanol precipitation. The concentration of purified sgRNAs was determined by measuring absorbance with a NanoDrop microvolume spectrophotometer.

#### Cas9 nickase digestion and linearization of supercoiled plasmids

Cas9 nickase (NEB M0650T) and sgRNA solutions were prepared according to NEB EnGen^®^ Spy Cas9 Nickase digestion protocol with 300 nM sgRNA and 30 nM Cas9 nickase. The reaction mixture was then allowed to incubate at room temperature for 10 minutes. Supercoiled plasmid DNA was then added to the reaction mixture at a final concentration of 3 nM. The reaction was then incubated at 37°C for 15 minutes after which it was placed on ice and digested with Proteinase K for 30 minutes. The reaction products were then phenol-chloroform ethanol precipitated and resuspended in a 10 mM Tris HCl buffer. Nicking efficiency was assayed via agarose gel electrophoresis by running the nicked product against an untreated supercoiled control. Reactions that showed greater than or equal to 95% nicked product as evidenced by the loss of mobility after conversion from a supercoiled to open-circle state were considered fully nicked and utilized for downstream assays. Samples were then digested with BsaI-HF®v2 (NEB R3733S) according to the manufacturer’s instructions. The reaction products were digested for 30 minutes at room temperature using Proteinase K (Roche, RPROTK-RO) and then were then purified via phenol-chloroform extraction followed by ethanol precipitation. The position of nicks and nicking efficiency was verified by Sanger sequencing using primers selected to hybridize to the nicked strand. A nick was confirmed through this method if the sequencing trace was terminated early at the expected site of the nick but successful in a complementary untreated control reaction. Sequencing was performed at the UC Davis Sanger sequencing core facility.

#### Cloning of pFC9ΔNt.BspQI series plasmids

pFC9 plasmid was digested with XbaI (NEB R0145S) and SapI-HF® (NEB R3156S) according to the manufacturer’s instructions to remove an existing Nt.BspQI recognition site. The large fragment resulting from this digest was then isolated via an agarose gel extraction. The plasmid ends were blunted using DNA Polymerase I, Large (Klenow) Fragment (NEB M0210) according to the manufacturer’s blunting protocol. The reaction products were isolated by phenol-chloroform extraction followed by ethanol precipitation and then ligated using T4 DNA ligase overnight at 16°C. The resulting ligated products were then transformed into DH5α competent cells. Single colonies were isolated, grown overnight, and plasmid was extracted via alkaline lysis followed by lithium chloride extraction and ethanol precipitation. The entire construct was then amplified with three independent sets of PCR primers using Q5 high fidelity DNA polymerase (NEB M0492S), each resulting in an insertion of an approximately 3.5 kb product with a Nt.BspQI site inserted at the forward primer binding site. The PCR products were then purified as previously described by agarose gel extraction and ligated using T4 ligase overnight at 16°C. The ligated products were then transformed into DH5α cells, grown up from single colonies, and plasmid DNA was isolated as previously described. The location and sequence of the Nt.BspQI insertions were verified via Sanger sequencing. Supercoiled plasmids were subsequently digested with Nt.BspQI (NEB R0644S) according to the manufacturer’s online protocol. Reaction products were purified using DNA binding spin columns and complete nicking of the substrate was verified in the same manner as Cas9 nickase-treated samples.

#### Estimation of R-loop junction free energy

The R-looper (https://github.com/chedinlab/rlooper_sim) source code was modified to allow for a zero junction energy value (a parameter) to be applied at a specific base pair range within a provided DNA sequence. Junction energy was then set to zero kcal/mol in a 20 bp region surrounding the location of Cas9 / Nt.BspQI induced nicks. A non-zero junction energy value was applied normally to all other base pair positions in the sequence. The modified R-looper was then run with increasing junction energy values for all other regions outside of the nick region. The position of predicted R-loops was then compared to experimental data to determine the junction energy which provided the best fit to the experimental data.

#### *In vitro transcription* with single strand binding protein

IVT reaction mixtures was prepared as previously described in *In vitro transcription reactions.* Co-SSB samples were supplemented with *E.coli* single-strand binding protein (8) (SSB; a kind gift from Dr. Stephen C. Kowalczykowski) to a final concentration of 2.5 μM before the addition of the appropriate polymerase. After termination of the transcription reaction, SSB was added to a final concentration of 2.5 μM to Post-SSB samples and allowed to incubate at 37° C for 20 minutes. Both Co and Post SSB samples were then digested with Proteinase K at 37° C for 30 minutes and then purified via phenol-chloroform extraction followed by ethanol precipitation. After the deproteinization and precipitation steps the samples were then bisulfite treated as described in *Non-denaturing bisulfite conversion*.

#### Non-denaturing bisulfite conversion

Non-denaturing bisulfite conversion of plasmid substrates was performed as previously described (9) using the EZ DNA Methylation-Lightning Kit (Zymogen D5030). Nicked samples were then treated overnight at 16°C with T4 DNA ligase (NEB M0202S) according to the manufacturer’s provided online protocol. This repair step ensures that the nicked strand can be PCR amplified, and R-loop signal can be subsequently measured.

#### SMRF-seq sample barcoding, PCR amplification, sequencing and analysis

Samples were amplified with a combination of forward and reverse primers that each incorporated a unique barcode. The combination of the forward and reverse primer barcodes uniquely identifies each sample within its specific flowcell. Primer binding sequences were designed in Primer3 (10) software and barcodes were generated using a custom Python script. Samples were amplified using Q5U® Hot Start High-Fidelity DNA Polymerase (NEB M0515S) for 25-30 cycles according to the manufacturer’s online protocol. 1-3 μl of bisulfite converted substrate was used for each reaction. In the case that the yield of one reaction was not sufficient, additional reactions using the original bisulfite converted sample were performed and then pooled. The target amplicon size was then purified via agarose gel extraction using BioBasic gel extraction kit (BS654). Sample concentration was quantified via agarose gel electrophoresis against 1kb linear DNA ladder (Thermo Fisher SM0311). SMRT cell PacBio libraries were constructed by the UC Davis DNA Technologies core according to manufacturer’s instructions and sequenced on a PacBio Sequel II instrument. Additional sequencing was performed at the California Institute for Quantitative Biosciences at UC Berkeley (QB3*-*Berkeley) on a PacBio Revio instrument. Fastq files were processed and R-loop footprints were called using the FootLoop program (https://github.com/srhartono/footLoop). R-loops were called using minimum parameters of 50 base pair minimum peak length (-t), at least 20 cytosines present in the putative R-loop (-w), and a minimum cytosine conversion rate of 35% (-t).

#### Assay for T7 initiation at nick sites

Supercoiled pFC9 was either nicked with Nt.BspQI or left untreated and digested with either a single digest of BamHI (NEB) or BamHI-HF and SacI-HF. The BamHI in this plasmid is located 5 bp upstream of the T7 promoter sequence while the SacI site is located 6 bp downstream. Therefore, digestion with only BamHI leaves the T7 promoter connected to the downstream R-loop forming SNRPN region while linearizing while digestion with both enzymes removes the T7 promoter from the larger plasmid as a ∼42 bp fragment. After digestion the large fragment of all digests were purified via agarose gel extraction and eluted into EB buffer. 400 ng of each sample was then used as input into an IVT. Each of the four samples were split into transcribed and un-transcribed aliquots and the IVT reaction using T7 RNAP was carried out as previously described. After the reaction the products were run on a 0.8% 1x TBE agarose gel for 2 hours at 60V. The gel was then post-stained with EtBr for 20 minutes and imaged.

#### *In vitro* transcription followed by S1 nuclease digestion

*In vitro* transcription reactions were performed as described. Samples were purified after termination of the transcription reaction via phenol-chloroform extraction followed EtOH precipitation. Samples were resuspended in 1x S1 nuclease buffer (Takara Bio 2410B) supplemented with MgCl_2_ to a final concentration of 6 mM to allow for simultaneous digestion with S1 nuclease (Takara Bio 2410B) and RNase H. RNase A was added to all samples at a final concentration of 5 ng/ μg. S1 nuclease was added to digested samples to a final concentration of 2 units / 100 ng of DNA and 3 units RNase H were added to digested samples. The reactions were kept on ice during preparation to avoid excessive digestion by S1 nuclease. All samples were brought to a final volume of 20 μl using PCR grade water and then incubated at 37°C for 30 minutes. After the incubation samples were removed and placed on ice and run on 1x TBE 0.8% agarose gel for two hours at 60V. The gel was then post-stained in EtBr for approximately 30 minutes, destained in DI water for 10 minutes, and then imaged using a BioRad GelDoc imager.

#### DNA secondary structure predictions

The complete sequence of pFC9 with the origin set at the PsiI recognition site to reflect sample digestion for AFM imaging was input into the RNAfold webserver (11) (http://rna.tbi.univie.ac.at/cgi-bin/RNAWebSuite/RNAfold.cgi) using the DNA parameters setting (12) and default parameters otherwise.

#### AFM sample preparation

For plasmid only samples, 10-15 ng of DNA was immobilized on a freshly cleaved mica disk in 20 μl of immobilization buffer (25 mM MgCl2, 10 mM Tris-HCl, pH 7.4) for 5 min. The mica was then washed 4 times with 20 μl imaging buffer (3 mM NiCl2, 20 mM HEPES, pH 7.4), before and a further 20 μl was added for imaging. For the S9.6 antibody binding experiment, 10 ng of DNA was pre-incubated with 1 ng of Anti-DNA-RNA Hybrid Antibody, clone S9.6 (Sigma Aldrich), and 7 µl of 1X DRIP buffer for 20 mins at room temperature. This reaction was then immobilized as previously in immobilization buffer (25 mM MgCl2, 10 mM Tris-HCl, pH 7.4) for 5 min, before washing and imaging in imaging buffer (3 mM NiCl2, 20 mM HEPES, pH 7.4).

#### AFM imaging

All AFM measurements were performed in liquid following a previously published protocol (13). All experiments were carried out in PeakForce Tapping imaging mode on either a Multimode 8 (Bruker) or FastScan Dimension XR AFM system (Bruker), using either PeakForce HRS-F-B (Bruker) or FastScan D (Bruker) probes. The PeakForce amplitude was set to 6-10 nm, the PeakForce Tapping frequency to 2-8 kHz and the PeakForce setpoints in the range: 7-15 mV, corresponding to peak forces of <70 pN. Various scan sizes were taken, maintaining a resolution of < 2 nm/px, at line rates of ∼1-5 Hz.

#### AFM image analysis

The freely available, open-source software TopoStats (14) was used to process the AFM data and analyze the DNA molecules (https://github.com/AFM-SPM/TopoStats). Briefly, the software loaded raw AFM images, carried out flattening, both line-by-line and plane flattening. Individual molecules were masked based on a height threshold to separate them from the background. A second flattening was carried out which excluded the grain, improving the flattening of the image. The height distribution of the flattened image was then shifted vertically to set the background to zero by calculating the mean of the non-grain containing data and subtracting that value from the image. Finally, a 1.1 px Gaussian filter was applied to reduce any high-gain noise. For classification and quantification of the R-loop features, the image analysis software FIJI was used to import the processed image files and manually trace the DNA strands. The features were classified as either “Blobs”, “Loops” or “Forks” depending on their morphology. The total contour length of the plasmids was measured as well as the distance to the feature and the length of the feature, if present.

#### Supplementary figures and figure legends

**Figure S1.**
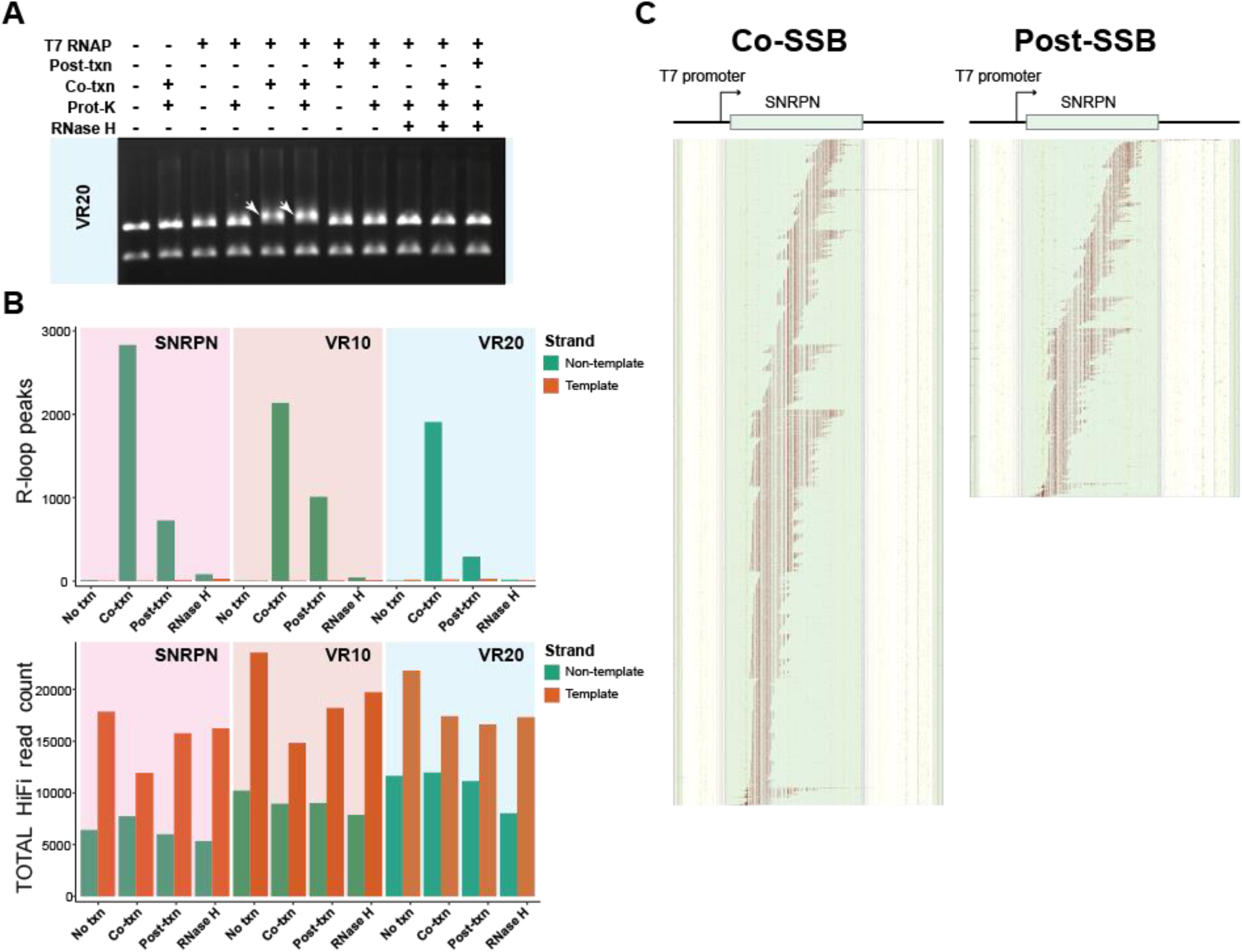
**A:** Agarose gel electrophoresis of the products of *in vitro* transcription reaction on the linearized VR20 plasmid under co and post SSB conditions. Upwards shifting indicating R-loop formation are indicated on lanes 5 and 6. Upward shifting is not reduced by Proteinase K treatment indicating this is not a result of a reduction of mobility due to SSB binding. This shifting is also sensitive to RNase H treatment (lanes 10 and 11). **B:** Barplot showing the number of measured R-loop peaks for all plasmids and SSB treatments utilized. **C:** Barplot of total number of reads for all plasmids and SSB treatments. **C:** R-loop footprints from *in vitro* transcription reactions of SNRPN containing plasmid under both co and post SSB conditions.

**Figure S2.**
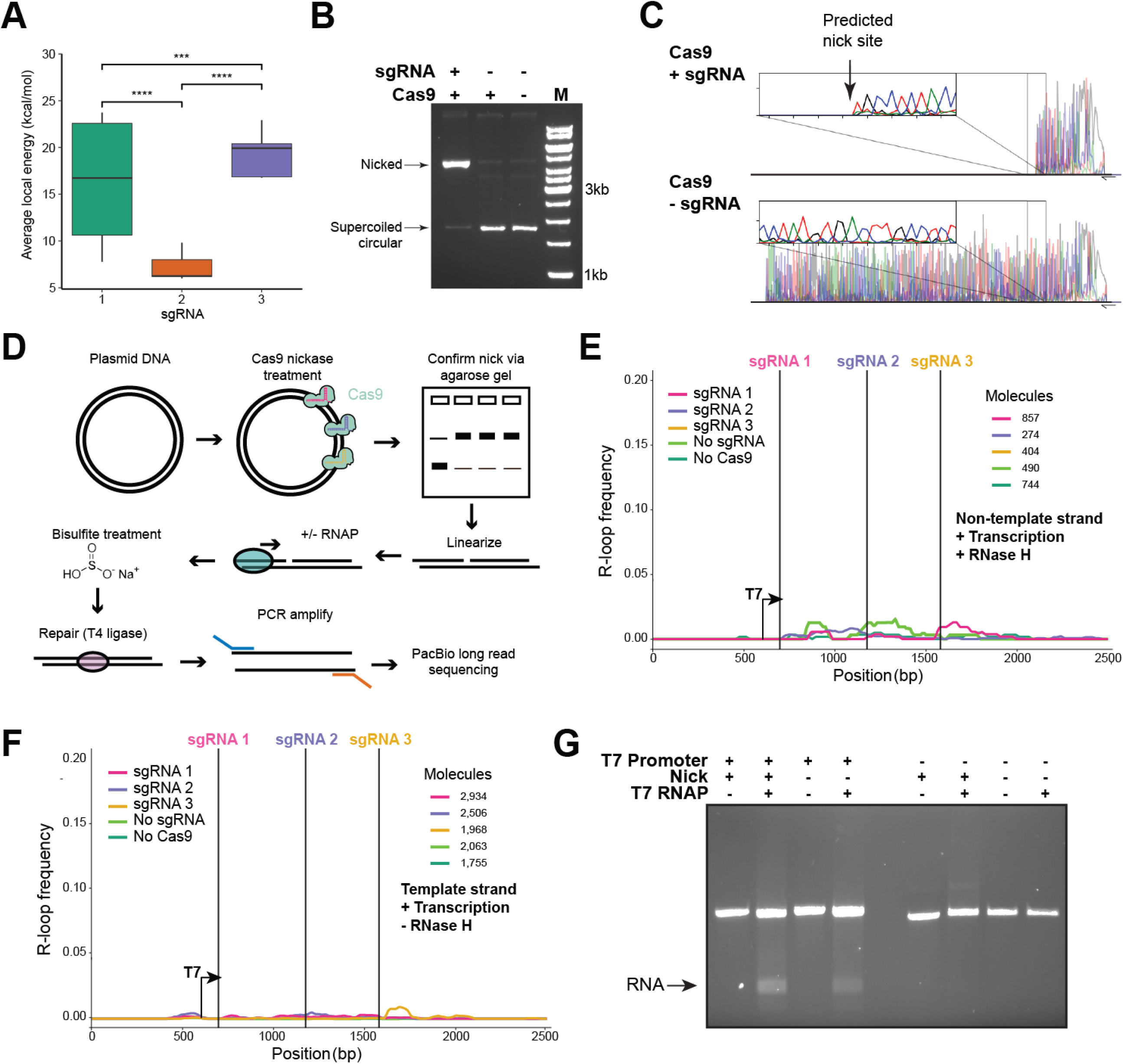
**A:** Boxplot showing local average energy in kcal / mol calculated using the R-looper model for the sequences within 50 bp of Cas9-induced nicks in the *SNRPN* region. **B:** Electrophoretic mobility shift assay comparing Cas9 and sgRNA-treated and untreated supercoiled DNA. **C**: Read traces from Sanger sequencing of a Cas9 and sgRNA-treated and untreated samples. The location of the predicted nick is highlighted. **D:** Diagram of the adapted SMRF-seq procedures for nicked substrates. See Sup Methods for additional details. **E**: R-loop frequency plot calculated from SMRF-seq data from the template strand of Cas9-nicked and unnicked plasmids. In this transcriptional context R-loop formation is expected on the non-template strand. **F**: R-loop frequency plot calculated from SMRF-seq data from transcribed and RNase H-treated Cas9-nicked and unnicked *SNRPN* containing plasmids on the R-loop-prone non-template strand. **G:** Agarose gel electrophoresis of the products of *in vitro* transcription reaction on linearized Cas9-nicked and unnicked plasmids containing, or not, the T7 promoter. RNA produced by transcription by T7 RNA polymerase can be seen as a smear at the bottom of the gel. Only plasmids containing the T7 promoter produced RNA regardless of their nicking status.

**Figure S3.**
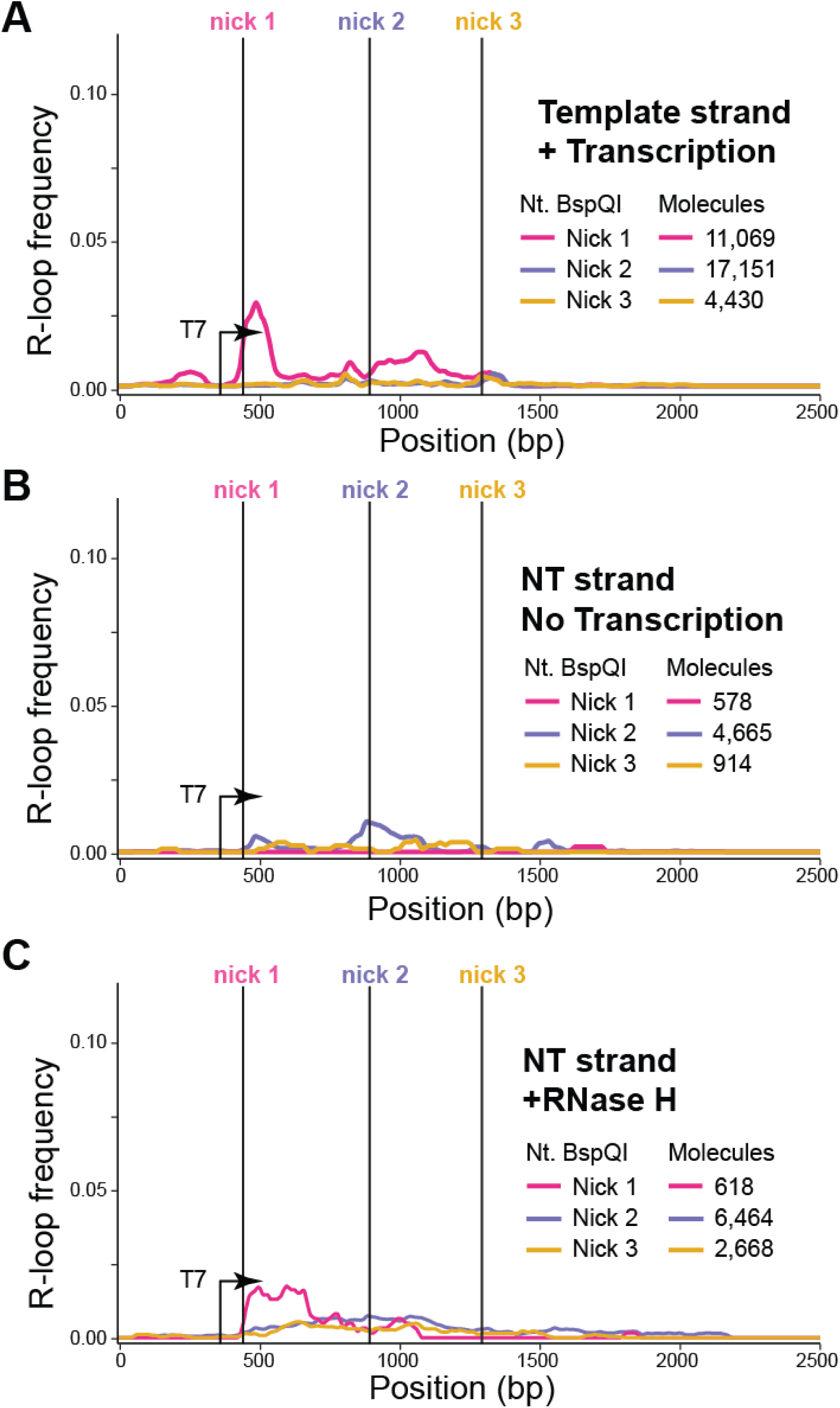
**A:** R-loop frequency plot from transcribed and Nt.BspQI-nicked plasmids calculated from SMRF-seq data on the template strand. **B:** R-loop frequency plot calculated from SMRF-seq data from untranscribed and Nt.BspQI-nicked substrates on the non-template strand. **C:** R-loop frequency plot calculated from SMRF-seq data from transcribed and RNase H1-treated Nt.BspQI-nicked substrates on the non-template strand.

**Figure S4.**
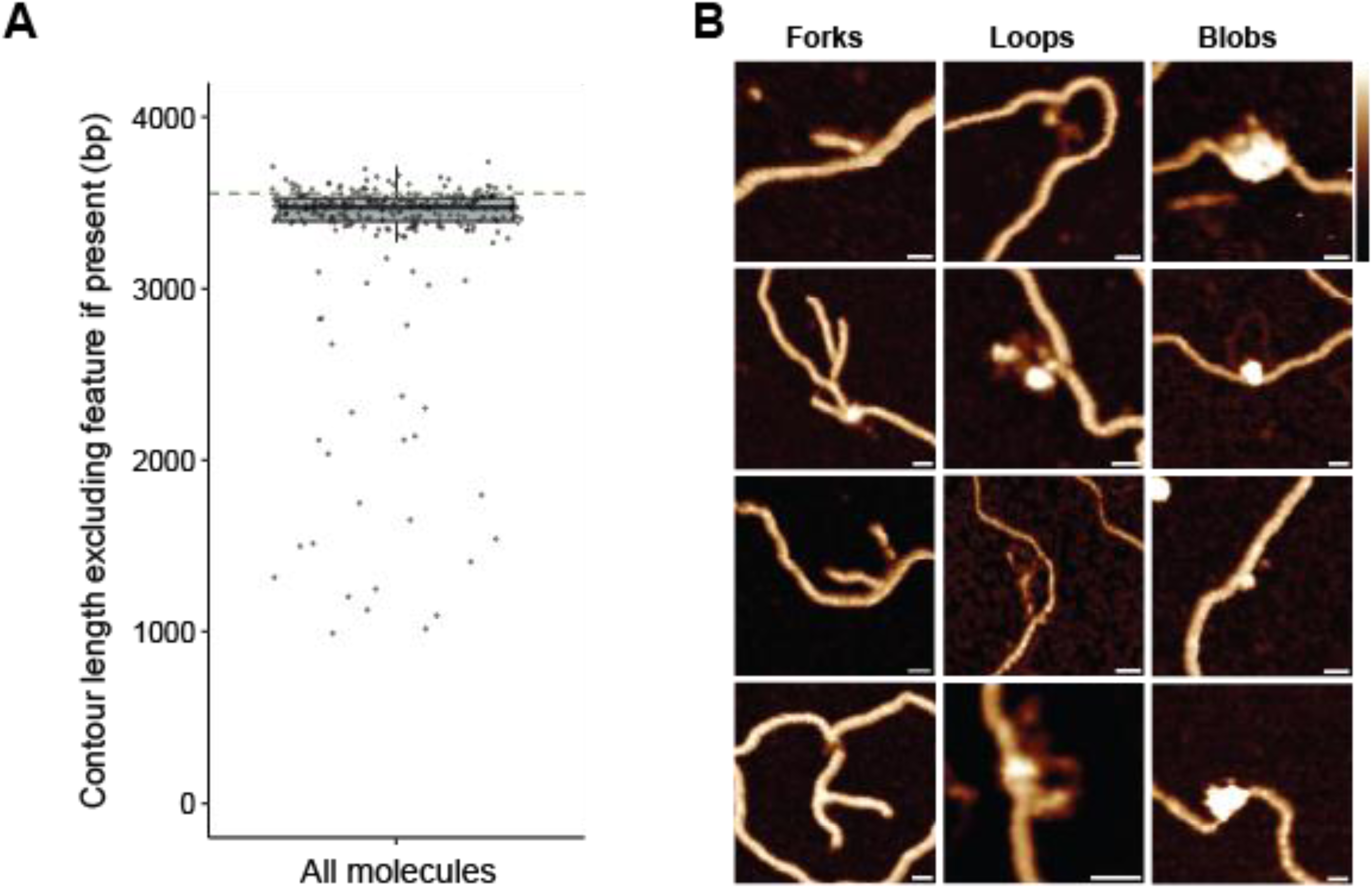
**A:** Boxplot showing distribution of molecule contour lengths measured via AFM in base pairs. The horizontal dashed green line shows the expected molecule size. **B:** Additional AFM images of the three classes of observed features; forks, loops and blobs. Scale bars indicate 10 nm. Height scale = -1 to 3 nm.

**Figure S5.**
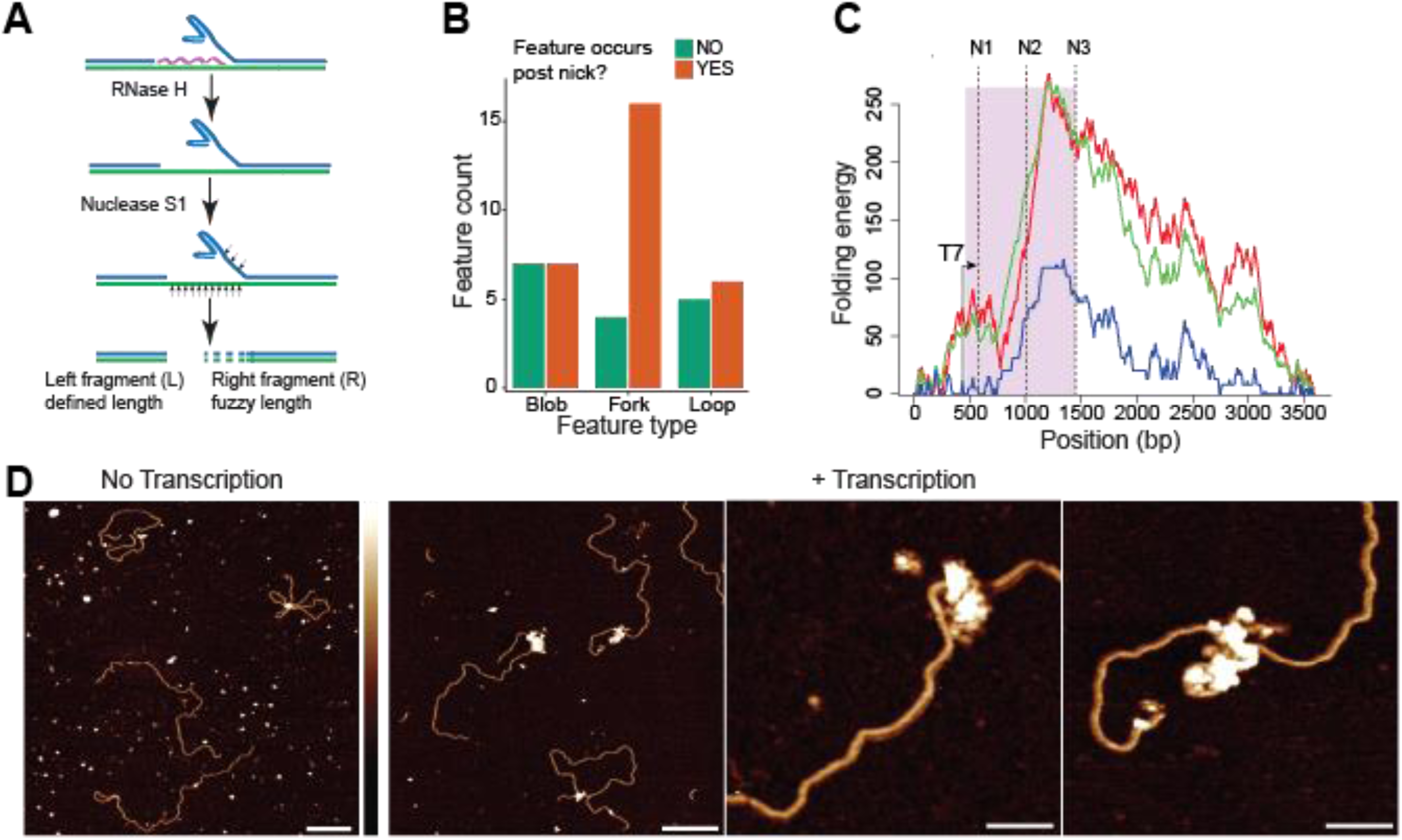
**A:** Cartoon diagram of model for digestion of a R-loop at a fork with Rnase H1 and S1 nuclease. **B:** Barplot showing the number of features, by feature type, that occur either before or after the site of a nick. **C:** DNA secondary structure favorability prediction over the SNRPN containing plasmid. The SNRPN region is highlighted in purple and the site of each of the three nicks are labeled and shown with dashed vertical lines. The location and direction of the T7 promoter is also shown. **D:** Representative images of transcribed and un-transcribed molecules treated with S9.6 antibody and imaged with AFM. The same concentration of S9.6 was used in both images. Scale bar for wide area images = 200 nm. Scale bar for close up images = 50 nm. Height scale = -1 to 3 nm.

**Figure S6:**
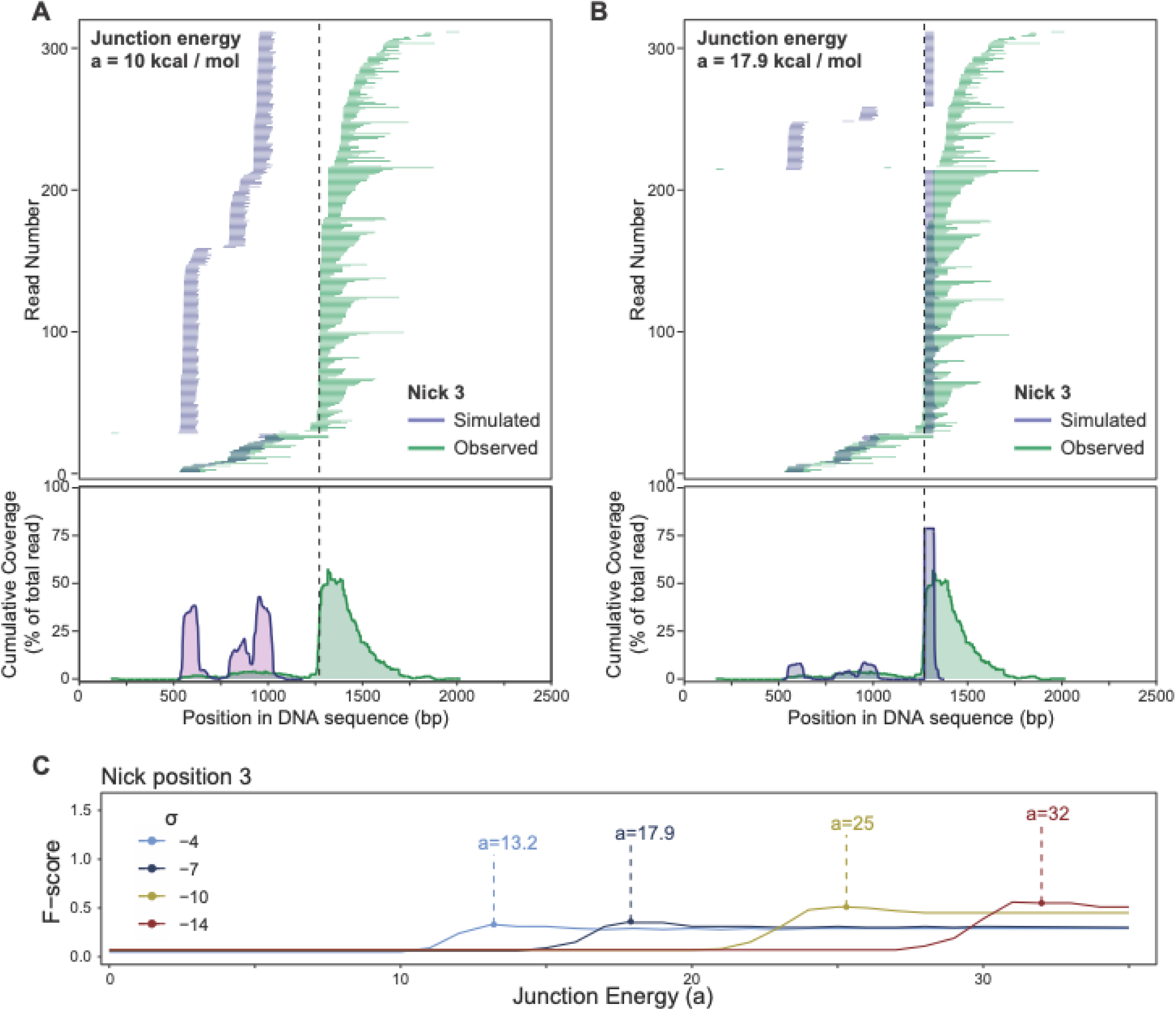
**A:** Observed (via SMRF-seq) R-loop locations from transcription of *SNRPN* containing plasmid nicked using Cas9 and sgRNA 3 vs. simulated R-loop locations from R-looper using the default junctional energy (a) value of 10 kcal / mol. The location of the nick is shown with a vertical dashed line. The top plot shows individual R-loop footprints while the bottom displays R-loop coverage. **B:** Identical to **A** but a junctional energy of 17.9 kcal / mol was utilized in the R-looper simulation. **C:** F-score plotted as a function of junction energy (a) for increasing levels of negative supercoiling (σ). The a value at the point with the highest f-score for each value of σ is shown.

